# A Drosophila eye modifier screen identifies TBC1D25 as a modulator of RAB21 phenotypes

**DOI:** 10.64898/2026.01.30.702430

**Authors:** Caroline Normandin, Sarah Dubois, Tomas Del Olmo, Steve Jean

## Abstract

Membrane trafficking is essential to maintain cellular homeostasis, enabling cells and organelles to exchange molecular components *via* vesicle transport. Therefore, it is tightly regulated, including by RAB GTPases. Among these, RAB21, which is primarily associated with early endosomes, plays a central role in coordinating endocytosis, sorting, and degradation. Like other RABs, it cycles between GTP- and GDP-bound forms. Although three specific guanine exchange factors (GEFs) for RAB21 have been identified, surprisingly, no GTPase-activating proteins (GAPs) have been found to directly modulate RAB21. Here, we describe a genetic modifier screen in *Drosophila* that identified Tre/Bub2/Cdc16 (TBC) domain family member 25 (TBC1D25) as a potential negative regulator of RAB21. We confirmed the RAB21–TBC1D25 interaction using co-immunoprecipitation and proximity ligation assays and further demonstrated that their association depends on the catalytic activity of TBC1D25. Genetic interaction studies revealed a functional link between TBC1D25 and RAB21 in autophagy and cargo sorting. Collectively, our results indicate that TBC1D25 negatively regulates RAB21, potentially by serving as a RAB21-specific GAP.

## INTRODUCTION

Membrane trafficking events, which can be defined as the vesicular transport of macromolecules between membrane compartments, play a crucial role in cellular functions and responses^1^. Trafficking events are dynamically modulated by the actions of a large array of proteins^2^, including those in the RAB GTPase family^3^. RAB GTPases represent the largest family of human small GTPases, with about 70 members^4,5^. They regulate multiple critical steps in vesicular transport, including cargo selection, vesicle formation, vesicle movement, tethering, and fusion^6,7^. Despite a few exceptions^8,9^, most RABs act as molecular switches that cycle between a GDP-bound inactive state and a GTP-bound active state^10,11^. RAB GTPase cycling is regulated by GEFs and GAPs^12^, which interact transiently with their target RABs and display varying degrees of specificity, with some affecting a single RAB and others modulating several^11,13^. GEFs promote a conformational change in RABs that enables GDP release, resulting in its replacement with GTP, which is present at high cytosolic concentration^14^. GTP binding alters the conformations of the switch I and II regions, allowing interactions between RABs and their effector proteins^10^. While their GEFs display wide structural diversity, RAB GAPs are almost exclusively TBC domain-containing proteins^10^. Once GTP hydrolysis is completed, RABs return to their inactive state. Multiple RAB GEF- and GAP-pairs have been described, but several RABs still lack identified GEFs or GAPs. RAB21 has well-described GEFs^14–16^, but no GAPs have been shown that act directly on it. Only TBC1D17 has been found to increase RAB21 GTP release *in vitro*^17^; however, in the same assay, TBC1D17 showed higher activity with RAB35 and also acted on RAB1, RAB5A, RAB5B, RAB5C, RAB8A, and RAB13^17^, arguing that TBC1D17 can target a wide range of RABs. More importantly, while TBC1D17 overexpression protects cells against Shiga toxin in a catalytically dependent manner, RAB21 depletion does not affect Shiga toxin trafficking, indicating independent functions^17^. Given this, it remains unclear if the *in vitro* activity of TBC1D17 translates into specific effects on RAB21 in cells, and as such, other GAPs could modulate RAB21 activity.

RAB21 plays important roles in various endosomal pathways, it modulates the endosomal recycling of various cargos in different contexts^18–23^, as well as multiple steps in autophagic flux^24–29^, with roles in autophagosome formation during starvation^28,29^ and in autophagosome-lysosome fusion in other contexts^24–26^. Its diverse biological functions include regulating neurogenesis^21,22^ and gut function^27^. In addition, RAB21 regulates the clathrin-independent endocytosis of β1-integrins in triple-negative breast cancer cells^30^, and processes it regulates have been identified as crucial in cancer metastasis^19,31^, treatment resistance^26^, and, more recently, in immuno-oncology^18^. Considering its diverse roles, identifying a RAB21 GAP would be a significant contribution to understanding its mechanisms of action.

In this study, we use a genetic modifier screen in the *Drosophila* eye to identify TBC1D25 as a modulator of RAB21. We confirm an interaction between RAB21 and TBC1D25 and demonstrate a functional link between the two proteins, showing that TBC1D25 affects two RAB21-mediated cellular processes in a catalytically dependent manner. These results indicate that TBC1D25 may represent a long-sought GAP for RAB21.

## RESULTS

### An eye modifier screen identifies TBC1D25, TBC1D13, and TBCK as potential RAB21 modifiers

To test for genetic interactions between Rab21 and GAPs, we leveraged its loss-of-function phenotype during *Drosophila* eye development **(Fig. 1A)**. We used the eye-specific NinaEGMR-Gal4^32,33^ to deplete Rab21 by RNAi. No eye phenotype was apparent when flies were kept at 18°C, as Gal4’s expression and activity are low at that temperature. However, dark patches, most likely corresponding to necrotic cells^34^, were apparent at 29°C, when Gal4 levels and activity are highest **(Fig. 1A, right panel)**. Given the high penetrance of the eye phenotype, we recombined NinaEGMR-Gal4 with the Rab21 RNAi construct to allow for the rapid screening of potential Rab21 modifiers **(Fig. 1B)**. We have previously validated the specificity of the IRRab21 line in different contexts^15,27,35^. Nonetheless, we tested whether we could rescue the necrotic eye phenotype by co-expressing upstream activating sequence (UAS)-driven Rab21 with the RNAi construct. Rab21 expression rescued the IRRab21 effect, indicating the possibility of identifying modifiers of Rab21 using this screen **(Fig. 1C)**. We reasoned that since RNAi-mediated depletion did not result in complete loss of Rab21, depleting a negative regulator could potentially restore activated Rab21 to levels sufficient to rescue the eye phenotype. We screened 26 TBC domain-containing proteins^36^, some with multiple RNAi lines when available, **(Fig. 1D)** and scored flies semi-quantitatively based on an organization index, with zero and three representing normal and completely necrotic eyes, respectively **(Fig. 1E)**. Depletion of TBC1D25, TBC1D13, and TBCK all resulted in improved eye structures and organization, suggesting that these proteins act as negative regulators of Rab21.

**Figure 1.**
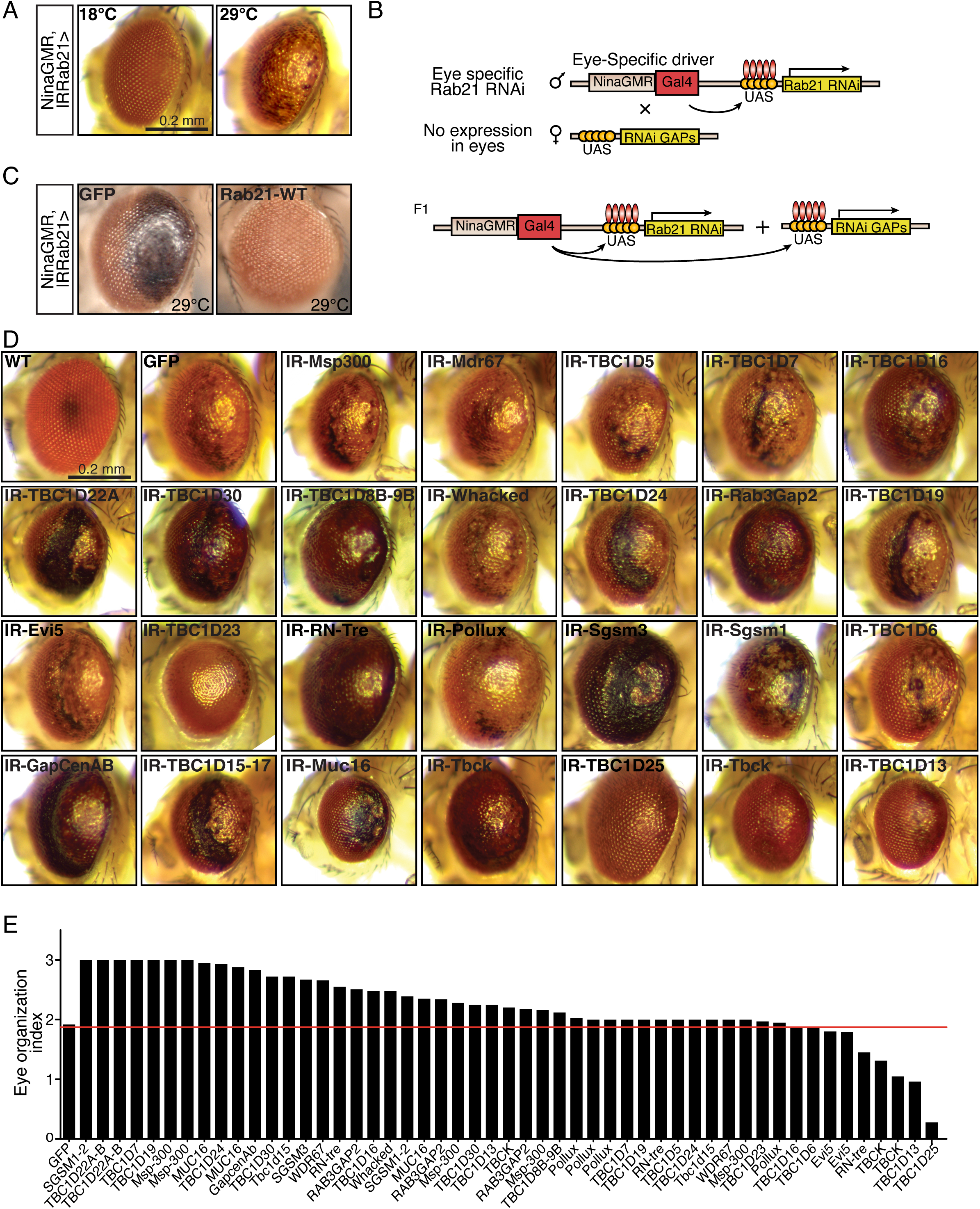
A RAB21 modifier screen identifies potential GAPs. **(A)** Representative images of adult *Drosophila* eyes expressing Rab21 RNAi constructs incubated at different temperatures. **(B)** Schematic of the modifier screen. NinaGMRGal4, UAS-IRRab21 flies were crossed with a collection of flies expressing UAS-driven RNAi constructs against fly GAPs. **(C)** UAS-Rab21-WT expression and not GFP expression rescues necrotic patches observed in Rab21-depleted eyes. **(D)** Representative images of adult *Drosophila* eyes coexpressing RNAi constructs targeting RAB21 and various GAPs. Some GAPs were tested using multiple RNAi lines; these have multiple scores. Flies were housed at 29°C until eclosion and their eye phenotypes were scored manually. **(E)** Eye organization index values upon Rab21 and GAP co-depletion. A normal eye phenotype was scored as 0, and a severity score was used to semi-quantitatively describe the phenotype: 1, eye disorganization with no necrotic patches; 2, necrotic patches covering < 50% of the eye surface; and 3, necrotic patches covering > 50% of the eye surface. The red line on the graph refers to the average score of Rab21-depleted eyes. Each bar graph represents the mean of at least 40 flies. No statistical analyses were performed, as the screen was conducted once.

### RAB21 and TBC1D25 interact in human cells

We moved to HeLa cells to assess potential protein interactions between human RAB21 and the three potential GAPs, since we have multiple functional RAB21 assays in these cells^20,25^. We performed coimmunoprecipitation studies in complete medium or after 30 min of full starvation, when our previous work revealed high RAB21 activity^25^. We also compared wild-type (WT), Q78L (GTP-bound), and T33N (GDP-bound) forms of RAB21 to assess if TBC1D25, TBC1D13, or TBCK would show nucleotide-preferential binding. We were unable to detect significant and consistent interactions with TBC1D13 or TBCK in any condition tested **(Suppl. Fig. 1A and B)**. However, TBC1D25 showed weak but reproducible binding with WT and GTP-locked RAB21 upon starvation **(Fig. 2A)**. We validated the interaction with endogenous TBC1D25 and green fluorescent protein (GFP)-RAB21, observing a weak interaction with WT RAB21 but no consistent binding with either GTP- or GDP-locked RAB21 **(Fig. 2B)**.

**Figure 2.**
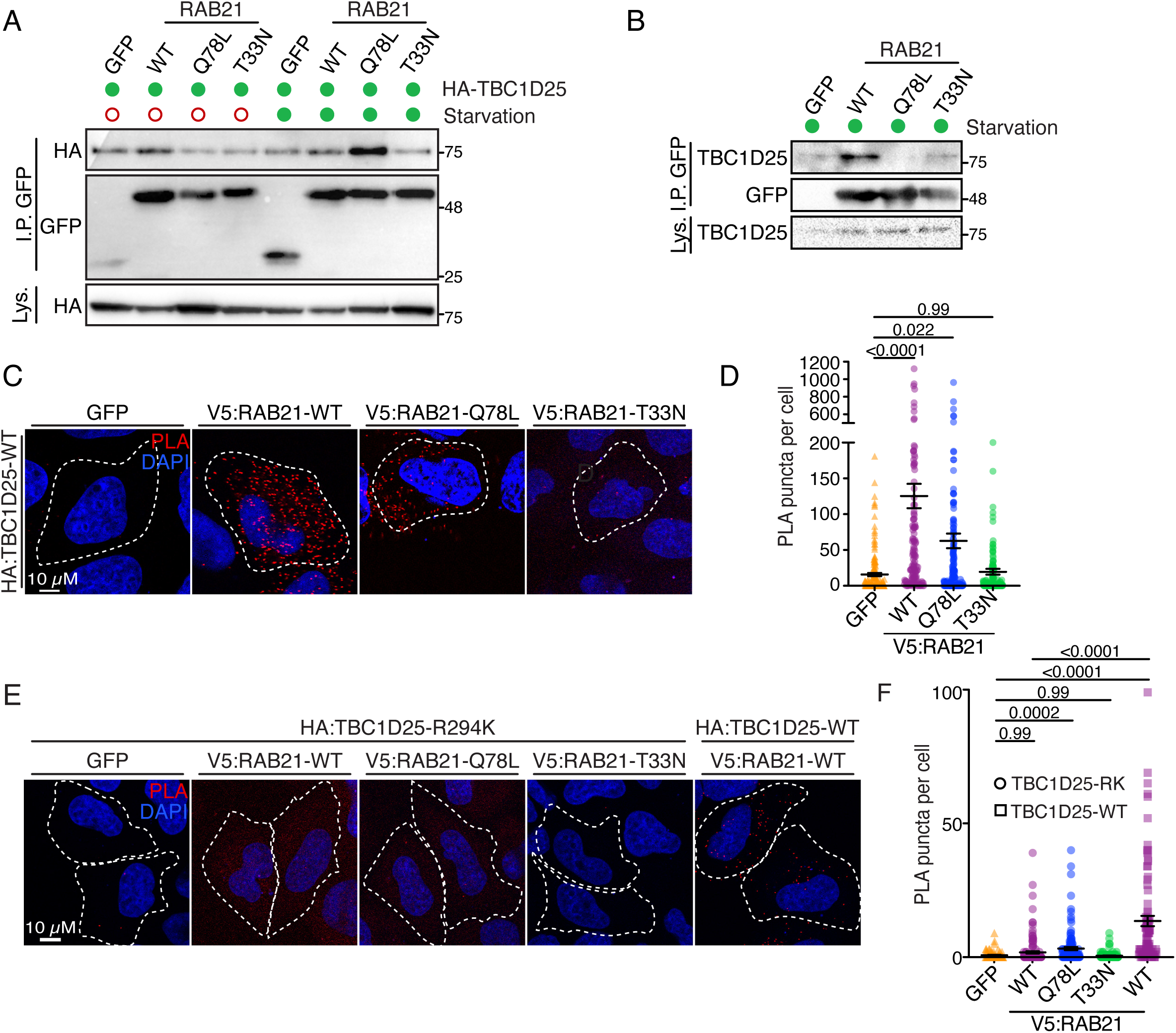
TBC1D25 interacts with RAB21 in HeLa cells. **(A)** Co-IP of GFP-tagged RAB21 WT, Q78L (GTP-locked), and T33N (GDP-locked) with HA-tagged TBC1D25 under normal growth conditions or after 30 min of starvation. Lysates were immunoprecipitated with anti-GFP antibodies and probed for HA to detect TBC1D25 binding. **(B)** Lysates from parental and GFP-RAB21 HeLa Flp-In T-REx cell lysates that were serum-starved for 30 min were subjected to GFP IP, then western blotted for GFP and endogenous TBC1D25. **(C)** PLAs of HA-TBC1D25 and V5-RAB21 constructs (WT, Q78L, and T33N) in normal growth conditions. **(D)** Quantification of the PLA puncta in (C). Individual points represent single cells, and the error bars are the SEM (*n*=3 independent experiments, ≥ 146 cells). **(E)** PLAs of HA-TBC1D25-R294K and V5-RAB21 (WT, Q78L, and T33N) in normal growth conditions. **(F)** Quantification of the PLA puncta in (E). Individual points represent single cells, and the error bars are the SEM (*n*=3 independent experiments, ≥ 90 cells). For all panels, *n* ≥ 3 independent experiments were performed. Dotted lines highlight transfected cells, which were detected by their GFP expression. In D and F, statistical significance was evaluated by one-way analysis of variance ANOVA (followed) by the Kruskal-Wallis *post hoc* test.

To further confirm the observed interaction, we performed proximity ligation assays (PLAs), which test for protein proximity and might detect more transient interactions^37^. We observed a high degree of close proximity between TBC1D25 and both WT and GTP-locked RAB21 **(Fig. 2C and D**). GAPs increase RAB GTPase activity through a conserved dual-finger mechanism^11^; therefore, we generated a catalytically inactive TBC1D25 variant (R294K)^38,39^. Interestingly, the proximity of TBC1D25 R294K to RAB21 was strongly reduced, with significantly fewer PLA puncta observed compared to WT TBC1D25 **(Fig. 2E and F**).

Given the genetic interaction in flies and the physical proximity in HeLa cells, we next tested whether TBC1D25 could act as a RAB21 GAP. Unfortunately, we were unable to establish *in vitro* GAP assays with bacterially purified TBC1D25 and either RAB21 or its known targets, RAB2A and RAB33B^38,39^ **(data not shown)**. We thus modeled binding between TBC1D25 and GTP-bound RAB2A, RAB33B, and RAB21 using AlphaFold3^40^. Interestingly, the interface predicted template modelling (ipTM) scores for RAB2A, RAB33B, and RAB21 were 0.86, 0.79, and 0.79, respectively **(Fig. 3A-C)**. These scores were close to the 0.8 score required for high-confidence prediction. Interestingly, ipTM scores dropped to 0.85, 0.75 and 0.73, respectively, when modeled with GDP, with RAB21 demonstrating the strongest decrease (not shown). Model confidence also decreased closer to the proximity surface with GDP. Significantly, all three RAB binding models included TBC1D25’s catalytic arginine (R294) and glutamine (Q332) residues forming hydrogen bonds with GTP **(Fig. 3D to F)**. This is consistent with its described mode of action and supports the potential for RAB GAP activity towards RAB21. Consistent with the PLA results, including the R294K mutation altered the predicted interaction surface with RAB21 by disrupting the contact between Gln332 and GTP, resulting in a decreased ipTM score of 0.69 **(Fig. 3G and H)**. Altogether, these AlphaFold3 structure predictions, combined with the genetic interaction screen and the co-IP and PLA results suggest a functional link between TBC1D25 and RAB21.

**Figure 3.**
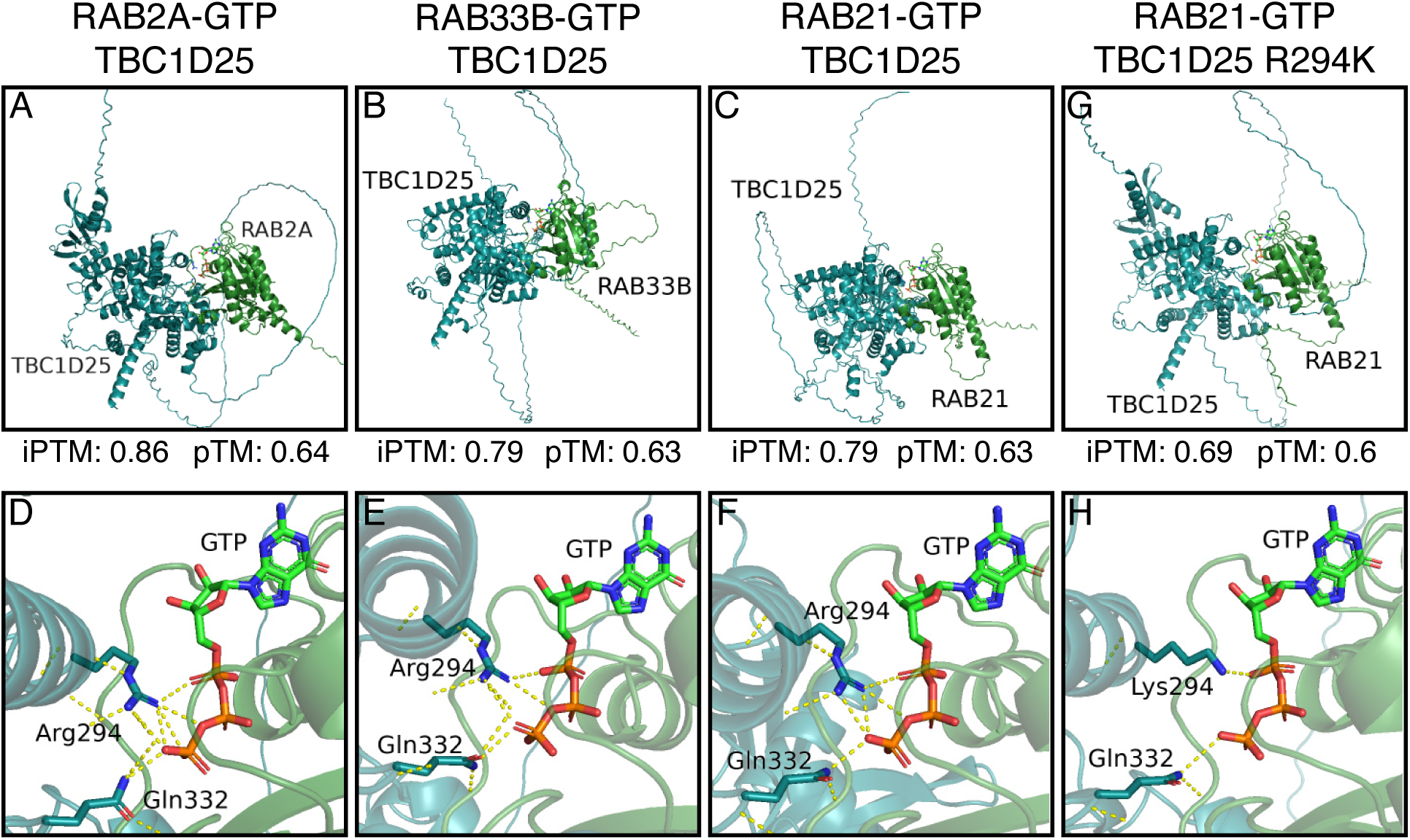
Structural modeling of TBC1D25 interacting with Rab GTPases reveals conserved binding interfaces and the impacts of Arg294 mutation (A–C, **G)** Predicted protein–protein interactions between WT TBC1D25 and GTP-bound forms of RAB2A (A), RAB33B (B), RAB21 (C), and with TBC1D25-R294K with RAB21 (G). Interaction confidence scores are indicated by the ipTM and predicted template modelling score (pTM) values. **(D–F, H)** Structural models highlighting the positions of TBC1D25’s catalytic Arg294 and Gln332 residues relative to RAB2A (D), RAB33B (E), RAB21 (F). (H) shows the positions of the lysine in the R294K mutant and Gln332.

### TBC1D25 overexpression decreases early endosomal RAB21 levels

RAB GAP overexpression is often associated with a decrease in the target RAB’s localization to its respective compartment^41^. Therefore, we reasoned that if TBC1D25 negatively regulated RAB21 in cells, TBC1D25 overexpression would affect RAB21 levels at early endosomes. We thus monitored GFP-RAB21 localization to early endosomes in TBC1D25 WT- or R294K-overexpressing cells. Consistent with a potential role for TBC1D25 as a RAB21 GAP, we observed decreased colocalization between RAB21 and early endosome antigen 1 (EEA1), an early endosomal marker **(Fig. 4A and B)**. Surprisingly, this effect was independent of TBC1D25 catalytic activity, and RAB21-EEA1 colocalization was also decreased in TBC1D25 R294K-overexpressing cells **(Fig. 4A and B)**, while no statistical differences in EEA1 puncta per cells were observed. In both TBC1D25 WT- and R294K-overexpressing cells, RAB21’s localization became more cytosolic. Altogether, these results imply that TBC1D25 overexpression affects RAB21 endosomal recruitment independently of its GAP activity.

**Figure 4:**
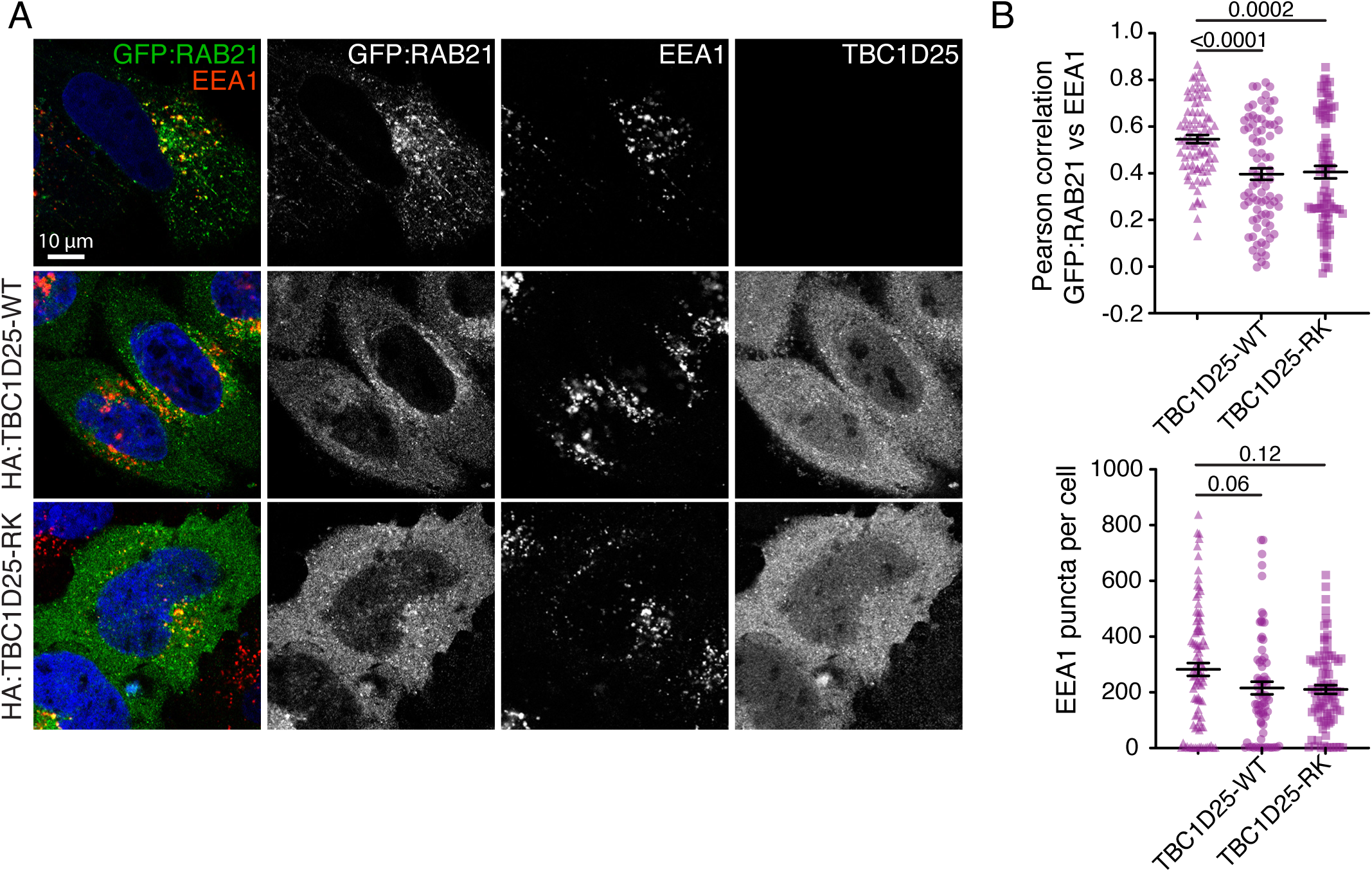
TBC1D25 modulates RAB21 localization to early endosomes. **(A)** Representative images of HeLa cells expressing WT or R294K TBC1D25. Cells were immunostained for EEA1 (red), HA (grey right-hand panels), and GFP (green). DNA was counterstained with DAPI (blue). **(B)** Quantification of GFP-RAB21–EEA1 colocalization (top) and the EEA1 puncta per cell (bottom). Individual points represent single cells, and the error bars are the SEM (*n*=3 independent experiments, ≥ 72 cells). Given the signal variability of EEA1, some cells display a total count of zero EEA1-positive vesicles, while they showed EEA1 vesicles. This strict thresholding was applied to ensure the most representative and reliable quantification of EEA1 structures across samples. Statistical significance was evaluated by one-way ANOVA followed by the Kruskal-Wallis *post hoc* test.

### TBC1D25 overexpression affects RAB21-linked processes

TBC1D25 overexpression has been shown to block autophagic flux, resulting in the accumulation of autophagosomes^39^. This work demonstrated that a RAB33B-TBC1D25 pathway is involved in autophagy, but the exact mechanism remains to be defined, and the authors suggested that TBC1D25 could also influence alternative pathways^39^. Our lab and others have shown that RAB21 depletion affects the late stages of autophagy at steady state^24–27^ and modulates autophagosome formation upon starvation^28,29^. Therefore, we asked whether the effect of TBC1D25 on autophagy is also partly mediated by RAB21. To test this, we overexpressed WT or R294K TBC1D25 in parental HeLa cells or a RAB21 knockout (KO) line **(Suppl. Figure 2A)** that we characterized previously^20,42^. WT TBC1D25 overexpression led to accumulation of the autophagosome marker microtubule associated protein 1 light chain 3 alpha (MAP1LC3A; hereafter referred to as LC3) in parental cells, while overexpressing the R294K mutant did not **(Fig. 5A and B)**, consistent with the findings of Itoh *et al*^39^. RAB21 KO cells displayed more autophagosomes than parental cells, and interestingly, overexpression of neither WT nor R294K TBC1D25 further increased their LC3 puncta **(Fig. 5A and B)**. These results suggest a model in which TBC1D25 affects autophagy partly through the negative regulation of RAB21, given the requirement for TBC1D25 catalytic activity.

**Figure 5:**
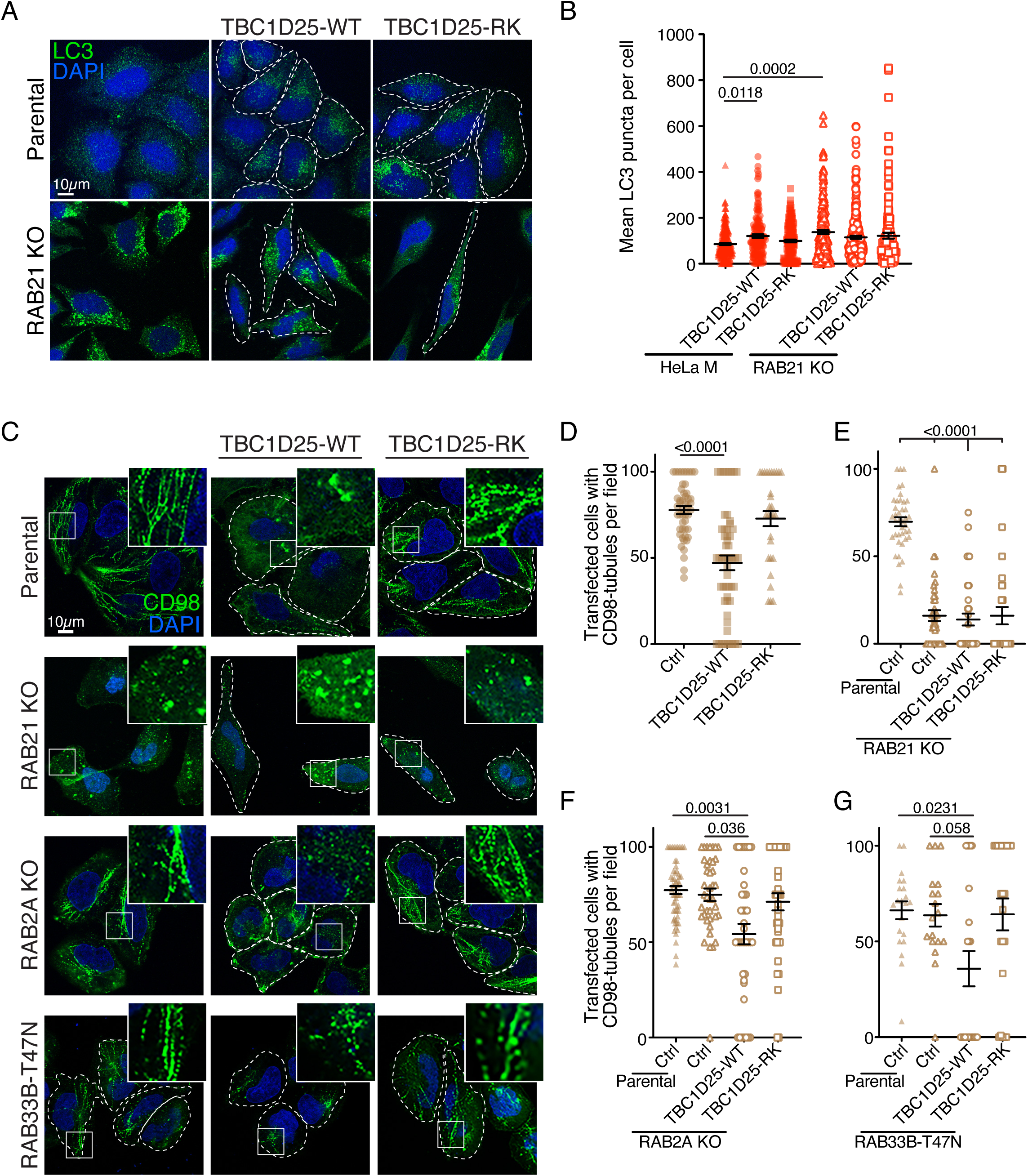
TBC1D25 regulates autophagic flux and CD98 trafficking in a RAB21-dependent manner. **(A)** Representative images of parental and RAB21 KO HeLa cells expressing WT or R294K TBC1D25. Cells were immunostained for LC3 (green) and counterstained with DAPI (blue). **(B)** Quantification of the LC3 puncta in (A). Individual points represent single cells, and the error bars are the SEM (*n*=3 independent experiments, ≥ 129 cells). **(C)** Antibody-uptake assays in parental, RAB21 KO, RAB2A KO cells, and RAB33-T33N-expressing HeLa cells overexpressing either WT or R294K TBC1D25. Cells highlighted by dotted lines represent transfected cells (TBC125 WT or TBC1D25 R294K or RAB33-T33N). Transfected cells, which were stained with HA (TBC1D25 or detected by their GFP fusion RAB33-T33N) are not displayed in a merge panel to simplify data representation. An acid wash was performed to remove any extracellular antibody before immunostaining for intracellular uptake. Insets display CD98-decorated endosomal tubules (*n*=3 independent experiments). **(D–G)** Percentages of cells displaying CD98-positive endosomal tubules per field in (D) parental, (E) RAB21 KO, (F) RAB2A KO, and (G) RAB33-T33N-overexpressing cells. Individual points represent single fields (averaging 6 cells per field), and the error bars are the SEM (*n*=3 independent experiments, ≥ 18 fields). In all graphs, statistical significance was evaluated by one-way ANOVA followed by the Kruskal-Wallis *post hoc* test.

To further investigate the connections between TBC1D25 and RAB21, we examined the effect of TBC1D25 overexpression on CD98 trafficking, which we have previously demonstrated to be modulated by RAB21^20^. In CD98 chase assays in parental HeLa cells^43^, overexpression of WT TBC1D25, but not the R294K mutant, significantly decreased the number of cells with CD98-labelled endosomal tubules **(Fig. 5C and D)**. This indicates that, as in RAB21 KO cells, WT TBC1D25 overexpression is sufficient to impair CD98 trafficking to endosomal-recycling tubules. However, RAB21 KO cells lacked CD98-decorated tubules, and this was unaffected by either WT or R294K TBC1D25 **(Fig. 5C and E)**. This is consistent with TBC1D25 negatively regulating RAB21 to affect CD98 trafficking. Finally, to define the impact of TBC1D25 on CD98 trafficking with respect to its known targets, RAB2A and RAB33B^38,39^, we generated a RAB2A KO population **(Suppl. Fig. 2B)**. As we were unable to generate a RAB33B KO line, we instead overexpressed a GDP-locked (T47N) dominant negative mutant. Neither RAB2A KO cells nor RAB33B dominant negative-expressing cells showed any defects in CD98 trafficking **(Fig. 5C, F, and G)**. Moreover, in the context of RAB2A KO or RAB33-T47N expression, WT TBC1D25 significantly decreased the number of cells harboring CD98 tubules, while the R294K mutant did not **(Fig. 5C, F, and G)**. These results suggest that TBC1D25-mediated modulation of CD98 trafficking is independent of its known RAB targets but is affected by RAB21 KO. TBC1D25’s effect on CD98 seems to be activity-dependent, providing further evidence that TBC1D25 may negatively regulate RAB21.

## Discussion

Multiple studies and methodologies have been used to define GAPs for specific RABs. Given the transient nature of these interactions, yeast two-hybrid or affinity purification approaches have often relied on the use of GTP-locked RABs^38,44–46^. Targeted phenotypic screens of GAPs have also been used to identify novel RAB-GAP pairs^17,36,41,47^. Altogether, GAPs for approximately 30 RAB have been described^48^. Here, we employed an eye modifier screen in the context of partial Rab21 depletion to identify GAPs whose depletion rescued the eye defects. We found three independent proteins whose depletion improved Rab21 depletion-induced eye defects, but we were only able to show an interaction with TBC1D25. TBC1D13 has been described as a RAB35 GAP^47^ and also interacts with RAB1 and RAB10 in yeast two-hybrid assays, although it did not act directly on these two RABs^47^. Given the role of TBC1D13 in trafficking^47^, the observed rescue could be an indirect effect of increased RAB35 activity, which may compensate for the loss of RAB21. TBCK knockdown also rescued Rab21 depletion, but we were unable to detect an interaction between the two proteins. The RAB targeted by TBCK is still undefined^49^, but it has recently been shown to be part of the FERRY complex^50,51^, which, among other proteins, contains RAB5^52^. TBCK loss of function has also been shown to decrease mTORC1 signaling in a context-dependent fashion^49,52^. Hence, it is conceivable that an increase in autophagy caused by mTORC1 inhibition could protect against cell death and prevent, to some degree, the appearance of necrotic patches in Rab21-depleted eyes. Even if TBC1D13 and TBCK do not act as RAB21 GAPs, their genetic interaction supports a potential functional link that could be worth exploring in specific contexts. Additionally, TBC1D4 and TBC1D32 (Pollux and Bromi in flies) were observed to interact with RAB21 in two recent studies^53,54^. While we did not test Bromi (TBC1D32), we did perform genetic interactions with *Pollux* and it did not strongly rescue Rab21 loss.

We demonstrated an interaction between RAB21 and TBC1D25 by co-IP and PLAs. However, we were unable to demonstrate direct GAP activity of TBC1D25 towards RAB21 *in vitro*, due to technical issues with the assay. It is conceivable that this assay failed because, as for TBCK and RAB5 in the FERRY complex, TBC1D25 and RAB21 require a protein scaffold to interact. While we cannot conclude whether TBC1D25 directly acts on RAB21 or indirectly through other RABs or processes, our modeling data support a potential GAP function, given the positions of arginine 294 and glutamine 332 in relation to GTP. In any case, our work reveals a functional relationship between RAB21 and TBC1D25—TBC1D25 overexpression affected both autophagic flux and CD98 trafficking in a RAB21-dependent fashion, suggesting that TBC1D25 regulates these processes partly through RAB21. The requirement for TBC1D25 catalytic activity in these processes also indicates that TBC1D25 acts as a GAP, either directly on RAB21 or on another RAB to indirectly regulate RAB21 activity. Unexpectedly, TBC1D25 modulated RAB21’s endosomal localization independently of its GAP activity. Although the RAB21 recruitment defects were phenotypically similar between WT and R294K TBC1D25, their molecular bases could differ—WT TBC1D25 overexpression could reduce RAB21-GTP levels, thereby precluding its association with early endosomes, while the R294K mutant could interact with specific RABs with greater affinity, modulating early endosome function and indirectly impacting RAB21 localization. Alternatively, if TBC1D25 acts as part of a larger complex, the R294K mutant could modulate complex composition, negatively affecting RAB21 recruitment independently of its GTP-bound status.

The physiological functions of TBC1D25 are mostly unknown, but it has been shown to regulate osteogenesis^55,56^. Given the functional link with RAB21 described here, it would be of interest to assess the roles of RAB21 in this context. The fly eye screen that revealed this genetic interaction may prove helpful in testing the functional relationships between other RABs and GAP proteins. Such screens could involve RNAi-based rescue of phenotypic effects, as done here, or overexpressing GAPs of interest and testing for rescue of resultant phenotypes with GTP-locked RAB expression. We hope that the identification of a functional link between RAB21 and TBC1D25 through a genetic modifier screen will help clarify functional relationships between other RAB-GAP pairs.

## MATERIAL AND METHODS

### Cell culture

Parental and HeLaM KO cells^20^ were cultured in Dulbecco’s modified Eagle medium (DMEM; Wisent, St-Jean-Baptiste, QC, CA) supplemented with 5% v/v fetal bovine serum (FBS; Wisent) and 0.01% penicillin/streptomycin (Wisent) at 37°C in 5% CO. Cells were starved by removing the full medium, washing twice with Earle’s Balanced Salt Solution (EBSS, Wisent), and incubating them in EBSS at 37°C in 5% CO_2_ for 30 minutes.

### Plasmids and cloning

TBC1D25, TBC1D13, and TBCK were cloned into the pcDNA3.1 3×HA plasmid to add an N-terminal triple HA tag. The cDNA of each GAP was amplified from HeLa cDNA by PCR. The pcDNA3.1 3×HA plasmid was digested with EcoRI, and each cDNA was inserted into it using the In-Fusion HD Cloning Kit (Takara Bio USA, Inc, San Jose, CA, USA.) according to the manufacturer’s instructions. The 3×HA-TBC1D25 R294K mutant was generated by site-directed mutagenesis using the Q5 Site-Directed Mutagenesis Kit (New England Biolabs Ltd, Whitby, ON, CA) according to the manufacturer’s instructions. The GDP-locked RAB33-T47N mutant in pEGFP-C1 was generated by Bio Basic (Bio Basic Inc, Markham, ON, CA) by gene synthesis. All plasmids generated in this study were verified by Sanger sequencing.

RAB2A KO cells were generated as previously done^20^. Briefly, three gRNAs against RAB2A (CCAATAGTAAGGTCATGCAC; TACATCATAATCGGCGACAC and CTGGCGGGCATCT TCTAACC) were cloned in pX330A2 and co-transfected with pEGFP-Puro (Addgene #45561) into cells at low passage numbers at a ratio of 15:1 with jetPRIME (Polyplus, New York, NY, USA). HeLaM cells (1×10) were plated in 6-well plates and transfected with 2 µg DNA total 24 h later using jetPRIME transfection reagent (Polyplus) following the manufacturer’s instructions. The DNA was added to 100 µL of jetPRIME buffer (Polyplus), then 4 µL of jetPRIME (Polyplus) was added and 50 µL in total added to 2 ml of full DMEM. The culture medium was replaced with fresh medium approximately 7 h after transfection. After 24 h of transfection, edited cells were selected for 48 h with 1 µg/mL puromycin (Wisent) in complete medium. RAB2A KO cells were validated by immunoblotting.

### Drosophila lines

The *Drosophila* line generated in this study was **[1]** *w[*];P{w[+mC]=GAL4ninaE.GMR}12, IR-Rab21*^109991^. Lines used for the screen were **[2]** *w;UAS-IR-Pollux,* VDRC:18319; **[3]** *w;UAS-IR-Pollux,* VDRC:104644; **[4]** *w;UAS-IR-Pollux,* VDRC:106969; **[5]** *w;UAS-IR-Pollux,* VDRC:109165; **[6]** *w;UAS-IR-Tbck,* VDRC:34780; **[7]** *w;UAS-IR-Tbck,* VDRC:108887; **[8]** *y[1] sc[*] v[1];UAS-IR-Tbck,* BDSC:57223; **[9]** *w;UAS-IR-Tbc1d23,* VDRC:40537; **[10]** *w;UAS-IR-Tbc1d23,* VDRC:110700; **[11]** *w;UAS-IR-Tbc1d16,* VDRC:22069; **[12]** *y[1] sc[*] v[1];UAS-IR-Tbc1d16,* VDRC:110067; **[13]** *w;UAS-IR-Whacked,* VDRC:22082; **[14]** *y[1] sc[*] v[1];UAS-IR-Tbc1d22A-B,* BDSC:32394; **[15]** *w;UAS-IR-Tbc1d22A-B,* VDRC:108659; **[16]** *w; UAS-IR-Tbc1d6,* VDRC:110561; **[17]** *w; UAS-IR-Tbc1d13,* VDRC:21001; **[18]** *w;UAS-IR-Tbc1d13,* VDRC:110396; **[19]** *w;UAS-IR-Tbc1d7,* VDRC:14705; **[20]** *w;UAS-IR-Tbc1d7,* VDRC:106667; **[21]** *w;UAS-IR-Rab3Gap2,* VDRC:27823; **[22]** *w;UAS-IR-Rab3Gap2,* VDRC:106905; **[23]** *y[1] sc[*]v[1];UAS-IR-Rab3Gap2,* BDSC:55328; **[24]** *y[1] sc[*] v[1];UAS-IR-GapcenAb,* BDSC:34976; **[25]** *y[1] sc[*] v[1];UAS-IR-Tbc1d8B-9B,* BDSC:32929; **[26]** *w;UAS-IR-Tbc1d19,* VDRC:25536; **[27]** *w;UAS-IR-Tbc1d19,* VDRC:100125; **[28]** *y[1] sc[*] v[1];UAS-IR-RN-tre.* BDSC:28670; **[29]** *w;UAS-IR-RN-tre,* VDRC:108670; **[30]** *w;UAS-IR-RN-Tre,* VDRC:28194; **[31]** *w;UAS-IR-Tbc1d25,* VDRC:108444 **[32]** *w;UAS-IR-Tbc1d5,* VDRC:24102; **[33]** *w;UAS-IR-Tbc1d24,* VDRC:44655; **[34]** *w;UAS-IR-Tbc1d24,* VDRC:108736; **[35]** *w;UAS-IR-Tbc1d15-17,* VDRC:20040; **[36]** *w;UAS-IR-Tbc1d15-17,* VDRC:109668; **[37]** *w;UAS-IR-Evi5,* VDRC:105146; **[38]** *y[1] sc[*] v[1];UAS-IR-Evi5,* BDSC:38350; **[39]** *y[1] sc[*] v[1];UAS-IR-Sgsm3,* BDSC:33729; **[40]** *w;UAS-IR-Wdr67,* VDRC:20316; **[41]** *w; UAS-IR-Wdr67,* VDRC:107134; **[42]** *w;UAS-IR-Sgsm1-2,* VDRC:48062; **[43]** *y[1] sc[*] v[1];UAS-IR-Sgsm1-2,* BDSC:28776; **[44]** *w;UAS-IR-Muc16,* VDRC:105591; **[45]** *y[1] sc[*] v[1]; UAS-IR-Muc16,* BDSC:44527; **[46]** *y[1] sc[*] v[1];UAS-IR-Muc16,* BDSC:53011; **[47]** *w;UAS-IR-Msp300,* VDRC:25906; **[48]** *w;UAS-IR-Msp300,* VDRC:105694; **[49]** *w;UAS-IR-Msp300,* VDRC:107183; **[50]** *y[1] sc[*] v[1];UAS-IR-Msp300,* BDSC:32377; **[51]** *y[1] sc[*] v[1];UAS-IR-Msp300,* BDSC:32848; **[52]** *w;UAS-IR-Tbc1d30,* VDRC:17314; **[53]** *w;UAS-IR-Tbc1d30,* VDRC:108779; **[54]** *w[*];P{w[+mC]=GAL4ninaE.GMR}12*.

### Coimmunoprecipitations

HeLaM cells were transfected as above with 2 µg DNA (1 µg of GFP-RAB21 (WT/Q78L/T33N) and 1 µg of HA-TBC1D25, HA-TBC1D13, or HA-TBCK). The following day, 48 h after cell seeding, HeLaM cells were starved in Earle’s Balanced Salt Solution (Wisent) for 30 min or kept in DMEM plus FBS and penicillin/streptomycin. Then, cells were washed twice in ice-cold 1× phosphate-buffered saline (PBS) and lysed in 300 µL of co-IP buffer (25 mM Tris-HCl, 1 mM ethylenediaminetetraacetic acid, 0.1 mM ethylenebis(oxyethylenenitrilo)tetraacetic acid, 5 mM MgCl, 150 mM NaCl, 2 mM Na VO, 10% glycerol, 1% NP-40, 1× protease inhibitor cocktail (Sigma-Aldrich, Oakville, ON, CA). Cells were scraped and incubated on ice for 20 min, then centrifuged for 10 min at 16,800 × *g* at 4°C. Supernatants were collected and transferred to new 1.5 mL tubes.

For co-IPs, GFP-Trap beads (Proteomics Platform, Université de Sherbrooke) were washed three times in co-IP buffer, non-specific binding sites were blocked with 1% bovine serum albumin (BSA; Wisent) in PBS for 1 h at 4°C on a nutator (Thermo Fisher Scientific, Whitby, ON, CA), and the beads were washed three more times with co-IP buffer. For each condition, 200 µL of supernatant was incubated with 10 µL of blocked GFP beads on a nutator at 4°C for 2 h. The beads were then washed four times with 1 mL of co-IP buffer and centrifuged for 30 s at 5,867 × *g*. After the final wash, the supernatant was removed, and 25 µL of 2× Laemmli buffer^57^ was added to the GFP beads. Beads were boiled for 5 min at 95°C before sodium dodecyl sulfate-polyacrylamide gel electrophoresis (SDS-PAGE).

Endogenous TBC1D25 co-IPs were performed on lysates from HeLa Flp-In T-REx GFP-RAB21 cells, which we generated previously^20^. GFP-RAB21 expression was induced by adding 10 ng/mL doxycycline to the medium for 24 h, then the cells were fixed with 0.5% formaldehyde for 15 min at room temperature (RT) with gentle agitation^58^. Glycine was added at a final concentration of 125 mM, and the cells were incubated for 5 min at RT. The cells were then washed twice with ice-cold PBS, and lysed and immunoprecipitated with GFP-Trap beads as above.

### Immunoblotting

Lysates and co-IPs were resolved by SDS-PAGE, adjusting the polyacrylamide percentage to the sizes of the proteins analyzed. Resolved proteins were transferred onto polyvinylidene fluoride (PVDF) membranes (Immobilon P, MilliporeSigma, Burlington, ON, CA) using a semi-dry Trans-Blot Turbo Transfer System (Bio-Rad Laboratories, Mississauga, ON, CA) for 10 min at 25 V and 2.5 A. After transfer, membranes were blocked in 5% milk (Carnation) in PBS with 0.1% Tween-20 (PBS-T) for 1 h with agitation at room temperature (RT). After blocking, primary antibodies were added in 5% milk in PBS-T, and the membranes were incubated overnight at 4°C on a nutator. Primary antibodies used in this study were anti-GFP (1:1000, SC-9996), anti-RAB2A (1:100, SC515612), and anti-HA (1:250, SC-393579) from Santa Cruz Biotechnologies (Santa Cruz, CA, USA), anti-HA (1:1000, 3724T) from Cell Signaling Technologies (Danvers, MA, USA), anti-TBC1D25 (1:500, HPA029197) and anti-tubulin (1:2000, T9026) from Sigma, and anti-RAB21 (1:1000, PA5-34404) from Invitrogen (Burlington, ON, CA). The next day, the membranes were washed four times in PBS-T for 5 min each, then incubated with secondary antibodies (horseradish peroxidase-conjugated anti-rabbit (111-035-144) and anti-mouse (115-035-146, both from Jackson ImmunoResearch, West Grove, PA, USA) at 1:10,000 in 1% milk in PBS-T for 1 h at RT. The membranes were washed four more times in PBS-T for 5 min each, then proteins were detected by enhanced chemiluminescence with either Luminata Forte (MilliporeSigma) or Clarity Max (Bio-Rad). A ChemiDoc MP system (Bio-Rad) or an Azure C280 (Azure Biosystems, Inc, Dublin, CA, USA) was used to visualize the protein signals.

### Proximity ligation assays

PLA was performed with the Duolink PLA kit following the manufacturer’s instructions (MilliporeSigma) with some modifications. Briefly, cells (3.0×10) were seeded on coverslips (Thermo Fisher, #1.5) in 24-well plates (Corning, Tewksbury, MA, USA). The next day, the cells were transfected with 500 ng of DNA as described above (100 ng of pEGFP-C1 to enable identification of transfected cells *via* GFP, 200 ng of either pcDNA-V5-RAB21-WT, Q78L, or T33N and 200 ng of an HA plasmid (pcDNA3.1-3×HA, pcDNA3.1-3×HA-TBC1D25 or 3×HA-TBC1D25-R294K). After 24 h, the cells were rinsed with PBS and fixed with 4% paraformaldehyde (PFA) in PBS for 15 min at RT. Cells were washed three times with PBS for 5 min each and then blocked with PBS containing 5% goat serum (Invitrogen) and 0.3% Triton X-100 (SigmaMillipore) for 1 h at 4°C. Primary antibodies (anti-V5, 1:1500, MilliporeSigma V8012 and anti-HA, 1:1500, Cell Signaling Technologies 3724T) were diluted in PBS containing 5% BSA and 0.3% Triton X-100 and applied to the cells overnight at 4°C. The next day, the cells were washed three times with PBS for 5 min each. Secondary antibodies (Duolink In Situ PLA Probe Anti-Mouse MINUS and Duolink In Situ PLA Probe Anti-Rabbit PLUS), were diluted in the PLA kit dilution buffer and incubated with the cells for 1 h at 37°C. After incubation, the cells were washed twice with buffer A for 5 min, then incubated with 2.5% ligase in 1× ligation solution for 30 min at 37°C. Cells were then washed twice with buffer A for 2 min each, then incubated with 1.25% v/v polymerase in 1× amplification solution for 100 min at 37°C. Finally, cells were washed twice with 1x buffer B for 10 min each and once with 0.01× buffer B for 1 min. Coverslips were mounted on slides using SlowFade Gold Antifade Mountant (Thermo Fisher Scientific) for analysis by confocal microscopy.

### Immunostaining and CD98 chase assays

Cells (3.0×10 HeLa cells or 4.5×10 RAB21-KO cells) were seeded on coverslips in 24-well plates and transfected 24 h later with 500 ng of either pcDNA3.1, HA-TBC1D25-WT, HA-TBC1D25-R294K, or GFP-RAB21-WT. The DNA was added to 50 µL jetPRIME buffer, 1 µL of jetPRIME was added, and 17.5 µL was added per well. The culture medium was replaced with fresh medium 7 h later. The next day, cells were washed once with PBS and fixed with 4% PFA for 15 min at RT. The PFA was removed, and the cells were washed three times with PBS for 5 min each then incubated in blocking buffer (PBS containing 5% goat serum and 0.3% Triton X-100) for 1 h at RT. Primary antibodies (anti-EEA1 (1:100, 3288T) and anti-LC3 A/B (1:200, 12741T) from Cell Signaling Technologies, and anti-HA (1:250, SC-393579) from Santa Cruz Biotechnologies) were diluted in PBS containing 10% w/v BSA and 0.3% Triton X-100 and applied to the cells overnight at 4°C. The next day, the cells were washed three times with PBS for 5 min each, then incubated with secondary antibodies for 1 h at RT (anti-mouse AlexaFluor 546 or 647, A11030 and A21236, respectively; anti-rabbit AlexaFluor 488 or 546, A11029 and A11035, respectively, all from Thermo Fisher Scientific), diluted at 1:250 in PBS containing 10% w/v BSA, 0.3% Triton X-100, and 1:10 000 4′,6-diamidino-2-phenylindole (DAPI)). Cells were then washed three times with PBS for 5 min each. Coverslips were mounted on slides using SlowFade Gold Antifade Mountant and analyzed by confocal microscopy.

For internalization assays, cells were seeded as above (4.5×10 RAB2A KO cells per well) and transfected as described above with 500 ng of either pcDNA3.1, HA-TBC1D25-WT, or HA-TBC1D25R294K, or 250 ng of pEGFP-C1-RAB33B-T47N and 250 ng of pcDNA3.1, HA-TBC1D25-WT, or HA-TBC1D25R294K. After 24 h of transfection, the cells were incubated with an anti-CD98 antibody (1:50, 315602 BioLegend) in full DMEM for 30 min at 37°C. Cells were then washed twice with ice-cold PBS, rinsed rapidly with 0.5% acetic acid and 0.5 M NaCl, pH 3.0, then rinsed twice with cold PBS for 5 min each. The cells were fixed with 2% formaldehyde (VWR) for 10 min at RT, washed twice with PBS for 5 min each, and blocked in PBS containing 10% goat serum and 0.1% saponin (MilliporeSigma) for 1 h at RT. The cells were incubated with an anti-HA antibody (3724T, Cell Signaling Technology), diluted to 1:500 in PBS containing 1% saponin and 10% goat serum, overnight at 4°C. The next day, the cells were washed three times with PBS for 5 min each, then incubated with anti-mouse Alexa Fluor 488 (A11034) and anti-rabbit Alexa Fluor 546 (A11035) secondary antibodies diluted to 1:250 in PBS containing 1% saponin, 10% goat serum, and 1:10,000 DAPI for 90 min at RT. The cells were washed four times with PBS for 5 min each, and the coverslips were mounted on slides using SlowFade Gold Antifade Mountant (Thermo Fisher). The presence of CD98-labeled recycling tubules was observed by confocal microscopy. In each image field, we calculated the percentage of cells overexpressing TBC1D25 or both TBC1D25 and RAB33B that contained CD98-labeled recycling tubules^20^.

### Confocal microscopy, image analysis, and figure generation

Immunofluorescence experiments were imaged on a Zeiss LSM 880 confocal microscope with a Plan-Apochromat 40×/1.4 oil objective and PLA experiments were imaged on an Olympus FV1000 confocal microscope with a Plan-Apochromat 63×/1.42 oil objective. Most channels were acquired sequentially to avoid bleed-through. Images were batch-exported in .tiff format using Zen Blue or Fluoview software.

After image acquisition, automated image analysis was performed using CellProfiler 3.0^59^, which enabled the identification of transfected cells and the study of desired parameters exclusively within them. Pipelines were created to identify GFP-transfected cells and PLA puncta for PLA experiments and to identify TBC1D25-positive cells for colocalization analysis and LC3 puncta counts. Once analyzed with CellProfiler, data were visualized using Prism (version 10, Dotmatics, Boston, MA, USA). CD98-positive tubules were manually quantified by counting the transfected cells with and without CD98 tubules in each field and calculating the percentage of cells with CD98 tubules.

All images were thresholded and cropped in Adobe Photoshop using linear parameters, with consistent thresholding within given experimental sets. Figures were assembled in Adobe Illustrator.

### Statistical Analyses

At least three independent replicates were performed for all experiments except the fly screen, which was performed once. In all quantifications except in Fig. 1D, data points represent individual cells or fields of view, the mean is indicated, and the error bars represent the standard error of the mean (± SEM). For all statistical analyses, a Shapiro-Wilk normality test was first conducted, and statistical tests were adjusted for parametric or non-parametric distributions.

Specific tests are described in the figure legends. When no statistical annotations are shown, it indicates that no significant difference was found. All statistical analyses were performed using Prism 10.

## Acknowledgments

We are grateful to laboratory members for their help throughout this work. We thank Dr. David Lambright for his suggestions on performing GAP assays and for sharing multiple protocols, Dr. Bruno Lemieux from the Protein Purification Core for his help with TBC1D25 purification, and High-Fidelity Science Communications for manuscript editing. Confocal imaging was performed in the Université de Sherbrooke photonic microscopy core. S.J. is a member of the Institut de Recherche sur le Cancer de l’Université de Sherbrooke (IRCUS) and the Fonds de Recherche du Québec Santé (FRQS)-funded Centre de Recherche du CHUS. This project was funded by a Canadian Institutes of Health Research grant (142305) and an institutional research chair from the Centre de Recherche Médicale de l’Université de Sherbrooke, both to S.J. S.J. is supported by a salary award from FRQS.

## Author contributions

This study was conceptualized by S.J. and C.N.. C.N. performed and analyzed the data for all the experiments, except Figure 2B, which was performed by T.D.O.. S.D. performed all the AlphaFold 3 modeling. C.N. and S.J. created the figures, performed statistical analyses, and wrote the manuscript, with feedback from T.D.O and S.D.. S.J. secured the funding for this research. All authors read and approved the final version of the manuscript.

