## Supplementary Figures for "A Drosophila eye modifier screen identifies TBC1D25 as a modulator of RAB21 phenotypes"

### SUPPLEMENTARY FIGURE 1

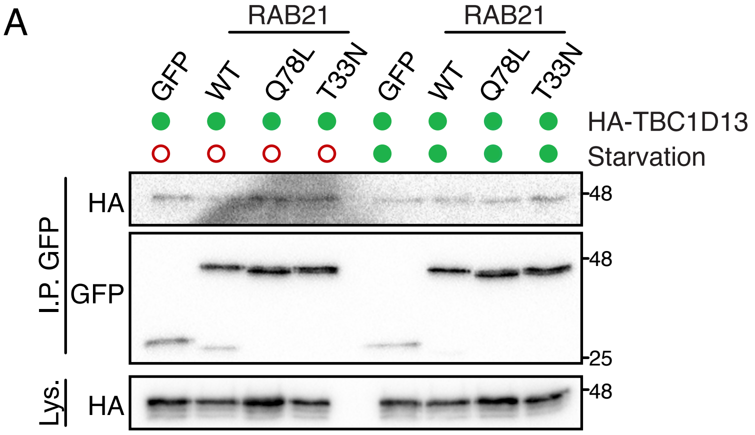


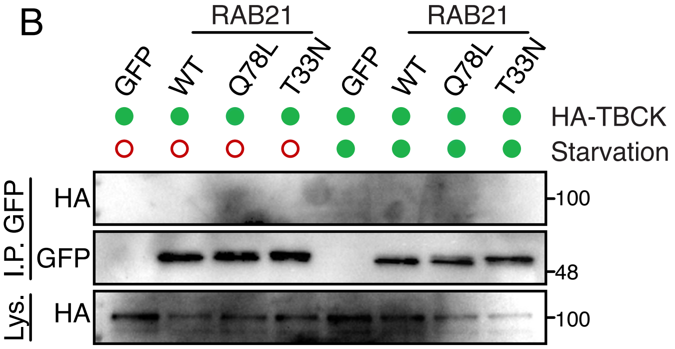


**Supplementary Figure 1: TBC1D13 and TBCK do not detectably interact with RAB21 by co-IP (A–B)** Lysates from cells expressing either GFP only or WT, Q78L (GTP-locked), and T33N (GDP-locked) GFP-RAB21 and **(A)** HA-TBC1D13 or **(B)** HA-TBCK that were grown under normal conditions or serum-starved for 30 min were subjected to GFP IP and western blotted for GFP and HA. Note that in B, GFP alone was not detected because it ran out of the 8% SDS-PAGE gel, given the molecular weight of TBCK.

### SUPPLEMENTARY FIGURE 2

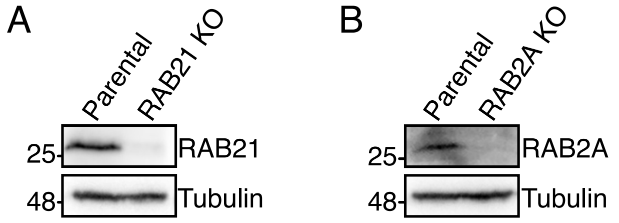


**Supplementary Figure 2: Validation of KO HeLa cell lines (A–B)** RAB21 and RAB2A expression levels in the **(A)** RAB21 KO and **(B)** RAB2A KO HeLa populations. Lysates were immunoblotted for endogenous RAB21, RAB2A, and tubulin (as a loading control).
